# Similar masking effects of natural backgrounds on detection performances in humans, macaques, and macaque-V1 population responses

**DOI:** 10.1101/2020.05.15.082537

**Authors:** Yoon Bai, Spencer Chen, Yuzhi Chen, Wilson S. Geisler, Eyal Seidemann

**Affiliations:** Center for Perceptual Systems, University of Texas, Austin, TX 78712, USA; Department of Psychology, University of Texas, Austin, TX 78712, USA; Department of Neuroscience, University of Texas, Austin, TX 78712, USA

## Abstract

Visual systems evolve to process the stimuli that arise in the organism’s natural environment and hence to fully understand the neural computations in the visual system it is important to measure behavioral and neural responses to natural visual stimuli. Here we measured psychometric and neurometric functions and thresholds in the macaque monkey for detection of a windowed sine-wave target in uniform backgrounds and in natural backgrounds of various contrasts. The neurometric functions and neurometric thresholds were obtained by near-optimal decoding of voltage-sensitive-dye-imaging (VSDI) responses at the retinotopic scale in primary visual cortex (V1). The results were compared with previous human psychophysical measurements made under the same conditions. We found that human and macaque behavioral thresholds followed the generalized Weber’s law as function of contrast, and that both the slopes and the intercepts of the threshold functions match each other up to a single scale factor. We also found that the neurometric thresholds followed the generalized Weber’s law and that the neurometric slopes and intercepts matched the behavioral slopes and intercepts up to a single scale factor. We conclude that human and macaque ability to detect targets in natural backgrounds are affected in the same way by background contrast, that these effects are consistent with population decoding at the retinotopic scale by down-stream circuits, and that the macaque monkey is an appropriate animal model for gaining an understanding of the neural mechanisms in humans for detecting targets in natural backgrounds. Finally, we discuss limitations of the current study and potential next steps.

**New & Noteworthy:** We measured macaque detection performance in natural images and compared their performance to the detection sensitivity of neurophysiological responses recorded in the primary visual cortex (V1), and to the performance of human subjects. We found that (i) human and macaque behavioral performances are in quantitative agreement, (ii) are consistent with near-optimal decoding of V1 population responses.

**Significance:** Natural selection guarantees that neural computations will be matched to the task-relevant natural stimuli in the organism’s environment, and thus it is crucial to measure behavioral and neural responses to natural stimuli. We measured the ability of macaque monkeys to detect targets in natural images and compared their performance to neurophysiological responses recorded in the macaque’s primary visual cortex (V1), and to the performance of humans under the same conditions. We found that (i) human and macaque behavioral performance are in quantitative agreement, (ii) are consistent with near-optimal population decoding of V1 neural responses, and (iii) that the macaque monkey is an appropriate animal model for gaining understanding of the neural mechanisms in humans for detecting targets in natural backgrounds.

## Introduction

The macaque monkey is an important animal model of the human visual system. The anatomy and physiology of their eyes, retinae, and early visual areas are similar to those of humans. Also, behavioral studies demonstrate similarity of human and macaque performance in visual detection and discrimination tasks, including similarity in their foveal contrast sensitivity functions (De Valois et al. 1974; Harwerth & Smith 1985a), spatial resolution across the visual field (Merrigan & Katz 1990), foveal wavelength discrimination functions (De Valois & Jacobs 1968), binocular summation (Harwerth & Smith 1985b), and binocular depth discrimination (Harwerth et al. 1995).

Recently there have been a number of studies measuring the human ability to detect targets in natural backgrounds (Bradley et al., 2014; Sebastian et al., 2017), which is critical for testing whether hypotheses derived from experiments with simple stimuli generalize to natural environments. These studies have shown that human thresholds follow a generalized Weber’s law:

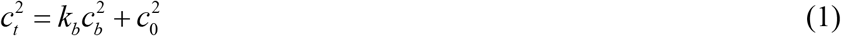

where *c_t_* is the root-mean-squared (RMS) contrast of the target at threshold, *c_b_* is the RMS contrast of the background, *k_b_* is the slope parameter, and 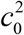the intercept parameter (*c*_0_ is the threshold on the uniform background).

However, there have been no studies quantitatively measuring macaque ability to detect targets in natural backgrounds. Here we take an initial step in this direction by comparing macaque and human psychometric functions for detecting a spatially-localized sinewave target in uniform backgrounds and in natural backgrounds of several different contrasts. In a separate experiment, we measured neural population responses in fixating monkeys to these stimuli in primary visual cortex (V1) using voltage sensitive dye imaging (VSDI), which provides a real-time measure of the local membrane potential responses over the entire region activated by the target stimulus. From these population responses we determined neurometric functions under the assumption of near optimal decoding at a coarse (retinotopic) spatial scale (Chen 2006; 2008).

We hypothesized that the neural processing underlying contrast masking effects is the same in humans and macaques, and that the generalized Weber’s law is the result of neural processing in the early visual pathway (retina and V1). If so, we expect equation (1) to hold, up to a single overall scale factor, for human behavior, macaque behavior, and macaque V1 population decoding. In other words, we predict that a single scale factor should align both the slopes and the intercepts of human and macaque threshold functions, and another single scale factor should align the slopes and intercepts of macaque and V1-neurometric threshold functions. This is a strong prediction because the slopes and intercepts need not vary together. However, we expect the overall scale factor (efficiency) to vary because of differences between humans and macaques in post-V1 processing (e.g., decision noise), and because of measurement noise in the V1 population responses.

We find that macaque and human behavioral thresholds measured under the same conditions are well-aligned by applying a single overall scale factor, and that macaque behavioral and V1-neurometric thresholds are well-aligned by applying a single overall scale factor. Thus, the results imply equivalent neural computations in humans and macaques, and that human and macaque thresholds can be explained by fixed-efficiency decoding of the retinotopic population responses in V1.

## Materials and Methods

### Detection task

We trained three macaques to detect a small horizontal Gabor target (σ=0.14°o, 4 cpd) in natural image backgrounds (yes-no detection task; Fig. 1C top). The natural image backgrounds were 4° in diameter, were windowed with a raised-cosine function (Figure 1B), and were centered on the location (2.5° eccentricity) where the target would appear when present. We used 50 of the same natural-image backgrounds used in a study measuring human detection thresholds (Bradley et al., 2014). The backgrounds were 512 pixels in diameter and were randomly sampled from ten large (4284 x 2844 pixels; 40 x 27 deg) calibrated images of natural scenes that contained no human-made objects (see Bradley et al., 2014 for details). The backgrounds were adjusted to have a Gaussian gray-scale histogram, and their contrasts were adjusted to 0% (uniform background), 1.875%, 3.75%, and 7.5% root-mean-squared (RMS) contrast (in one of the monkeys we only tested background contrasts of 0%, 3.75%, and 7.5%). Background mean luminance was set to the average luminance of the uniform grey background (30 cd/m^2^). For each background contrast, we structured trials into blocks such that for each block the contrast of the Gabor target was fixed and the target was present on 50% of the trials. For each block, we randomly chose the target contrast from a list of fixed values that ranged from 0.74% to 7.1% RMS contrast (3.125% to 30% Michelson contrast).

**Figure 1.**
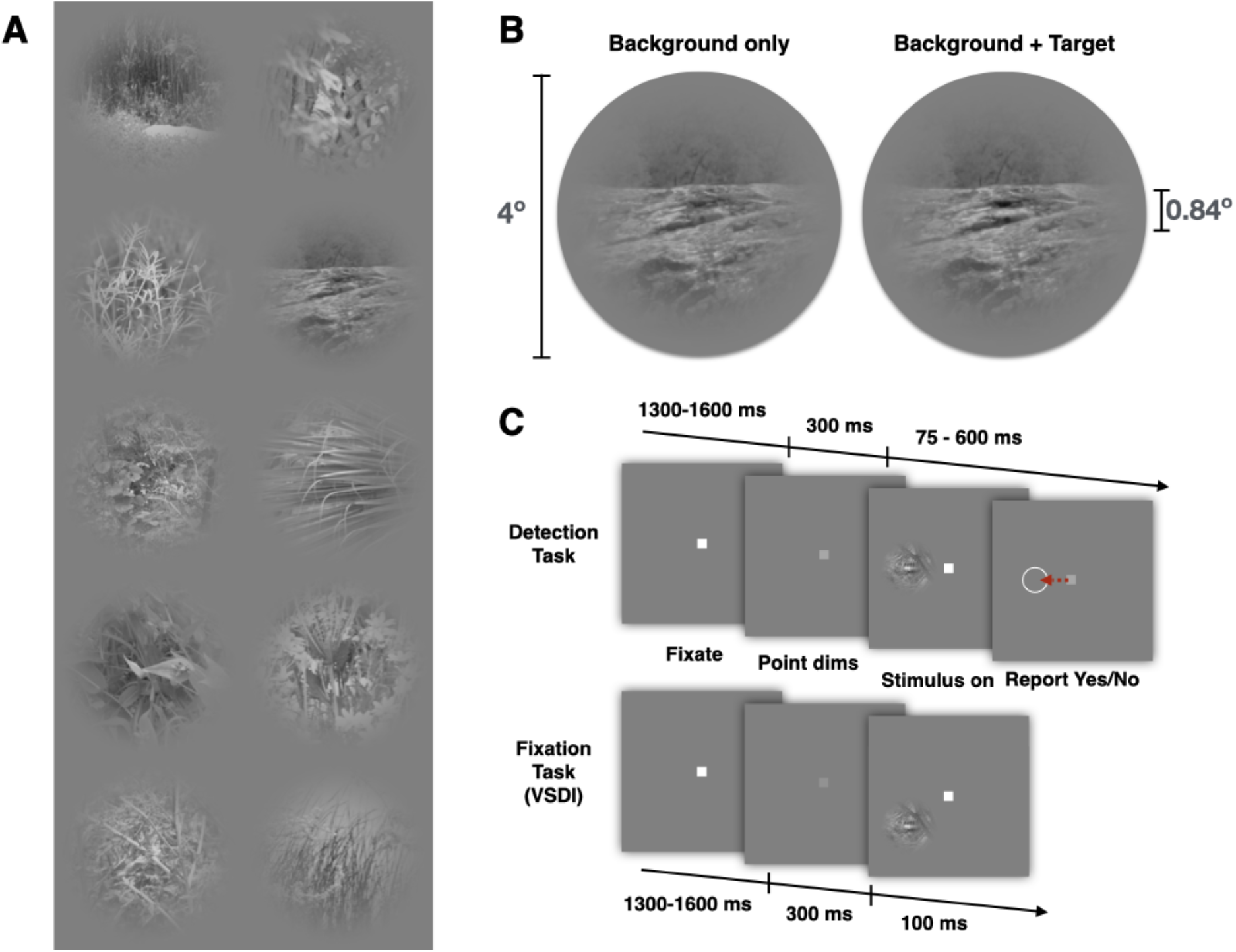
Target detection task. **(A)** Each natural-background patch was cropped and windowed from one of the natural scene images used in Bradley et al. (2014). In the behavioral experiments 50 backgrounds were used. Shown here are the 10 backgrounds used in the VSDI experiments, which were picked to be diverse and representative. **(B)** Background patches were 4° in diameter. The target was a horizontal Gabor (σ=0.14°, 4 cpd). In this example, a low-contrast target is added to a background with an RMS contrast of 15%. **(C)** Detection and fixation tasks. Detection performance was measured in a single-interval forced-choice task. Each trial began with a brief audible cue and a fixation point that was centered on a uniform grey background. The monkey was required to initially maintain fixation. Following this period, the fixation point dimmed and the stimulus was presented at 2.5° eccentricity. For target-absent trials, the monkey was required to maintain fixation to receive a juice reward. For target-present trials, the monkey was required to saccade to the target location and maintain gaze at the target location for an additional 200 *ms* to receive a reward. The stimulus was turned off as soon as gaze left the fixation window and a circle cue was presented to help maintain fixation at the target location. The report time window was 75 - 600 *ms* following stimulus onset. In the fixation task, the monkey was required to maintain fixation during the entire trial.

Each trial began with a brief audible cue and a fixation point that was centered on a uniform grey background displayed on a CRT monitor positioned at a distance of 108 cm. The monkey was required to maintain fixation for 1300-1600 ms to ensure that it was fully engaged in the task. Following this period, the fixation point dimmed for 300 ms and then the stimulus was presented at a parafoveal location (2.5° eccentricity). The stimulus consisted of a natural background either with or without the Gabor target. In target-absent trials, the monkey was required to maintain fixation for an additional 1 second to receive a juice reward. Trials were categorized as “false-alarms” when the monkey shifted gaze to the center of the background region. In target-present trials, the monkey was required to saccade to the target location and maintain gaze at the target location for an additional 200 ms to receive reward. The report time window was 75 ms to 600 ms after stimulus onset. Target-present trials were categorized as “misses” when the monkey held fixation beyond the report time window. In relatively few cases (< 5%), we aborted the trial when the monkey did not hold gaze precisely (1 ° diameter fixation window) or when the monkey made a saccade to an arbitrary location. These aborted trials were repeated at the end of the block. Across target-present trials, the monkeys’ median reaction times were approximately 200 ms. The visual stimulus was presented for up to 300 ms, but was switched to uniform gray as soon as the monkey initiated a saccade. Target-present and target-absent trials were randomly interleaved and the duration of each trial lasted from 2 to 3 seconds, with a fixed inter-trial interval of 2.5 seconds for correct responses. Audible feedback was given at the end of each trial. For correct trials (hits and correct-rejections), a juice reward accompanied a positive audible feedback. For misses and false-alarms, an extra 3 second intertrial interval was added with a negative audible feedback (for a fixed experiment duration, incorrect trials reduce the total amount of juice reward).

The number of experimental sessions for the three monkeys were 16, 19, and 16. In each session two of the four background contrasts were tested—five blocks of trials for one background contrast followed by five blocks of trials for the second background contrast. Each block of trials was for a different randomly-selected (without replacement) target contrast. The number of trials in a block was 100 (each background was presented once with target present and once with target absent).

### Estimating behavioral detection performance

Psychometric functions were estimated separately for each monkey and background contrast. To take into account the effect of criterion bias, we calculated detectability (d’) values for each stimulus condition using the standard formula,

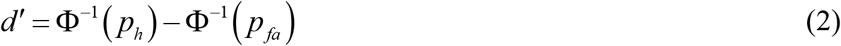

where *p_h_* and *p_fa_* are the proportion of hits and false alarms. For each monkey and each stimulus condition, the *d’* values were averaged across experiment sessions and converted to an effective (maximum) percent correct:

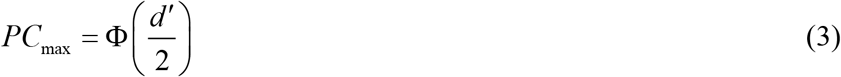

These accuracy values were then fit with a descriptive function. This descriptive function was the UNI (uncertain normal integral) function, which is similar to the familiar Weibull function but more principled (see Geisler 2018),

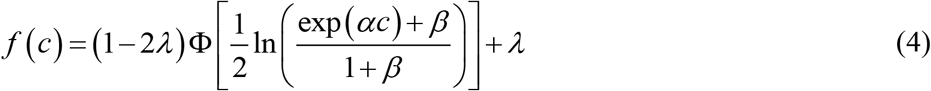

where *c* is the RMS contrast of the target, *α* is a parameter that is dependent on the background contrast, *β* is a parameter that varies with the level of intrinsic position uncertainty of the target, and *λ* is the lapse rate. In this study, *β*was fit as a constant across all conditions, because intrinsic position uncertainty varies with retinal location (Michel & Geisler 2011) and the target (when present) was always positioned at the same retinal location. Lapse rate was also fixed for a given subject (or when fitting the average across subjects). Thresholds were defined to be the target contrast giving a percent correct (PC_max_) of 69% (*d’* = 1).

### VSD Imaging

Wide-field imaging with voltage-sensitive dyes was used to record neural population activity at a high resolution in space and time (Shoham et al., 1999). Before each imaging experiment, voltage-sensitive dyes (RH 1691 or RH 1838) were topically applied to the cortex through a surgically implanted chamber. Measurements were made after an approximately 2 hour waiting period, which allowed the VSD molecules to bind with neural membranes. Fluorescence from neural activity was recorded using a commercial imaging system (Optical Imaging, Inc.). The imaging system was configured to record from a cortical region of approximately 8 x 8 mm, capturing V1 population responses over the whole region where activity is elicited by the Gabor target. Imaging data were collected at 100 Hz or 110 Hz where each frame was 512 x 512 pixels. VSD molecules were excited by light at 630 nm. Fluorescence signals were measured through a dichroic mirror (650 nm long-pass filter) and an emission filter (RG 665). Stimulus presentation and data acquisition was synchronized with the monkey’s EKG signal to minimize trial-to-trial variations due to cortical pulsations from the heartbeat. VSD responses are a linear function of the locally integrated sub-threshold neural activity from dendrites and axons in the superficial layers of the cortex (Grinvald and Hildesheim, 2004; Chen 2012). More details about optical imaging with VSD in behaving monkeys are described elsewhere (Seidemann et al., 2002; Slovin et al., 2002; Arieli et al., 2002).

### VSDI fixation task

VSDI measurements were made in two monkeys (different from those in the detection task) trained to perform a fixation task (Fig. 1C bottom). Each imaging trial began with an audible cue and a small fixation point presented on a uniform blank screen. The monkey was required to first hold fixation for 1000 to 1300 ms. Following this “initial” phase, the fixation point dimmed to indicate the beginning of the “stimulus” phase of the trial. The monkey did not receive a juice reward if fixation was broken during either the initial or stimulus phase of the trial.

The stimulus conditions in the fixation task were essentially the same as in the detection task. The background luminance was fixed at 30 cd/mm^2^, and the RMS contrasts of the backgrounds were 0%, 1.875%, 3.75%, and 7.5% for monkey 1, and 0%, 3.75%, and 7.5% for monkey 2. For each level of background contrast we repeated five levels of target contrasts in a random fashion (Gabor target contrast (RMS): 0%, 0.74%, 1.5%, 3%, 6%, and 9.5%; Michelson contrast: 3.125%, 6.25%, 12.5%, 25%, and 40%)

In each session, responses were measured for a 0% (uniform) background and for one of the higher contrast backgrounds.

Although the stimulus conditions were essentially the same as in the detection experiment, only a subset of ten natural background patches were tested. For each background contrast, each of the 10 background patches was presented once without the target and once with the target at each contrast level. Fewer backgrounds were tested because less time was available for making the VSDI measurements than the behavioral measurements. The natural background patches were manually selected to include a range of representative spatial structures from the full set (e.g., dense, sparse, oblique, horizontal, and vertical structure; see Figure 1A).

Stimulus presentation was somewhat different for the two monkeys. For monkey 1 the stimulus duration was 100 ms, and for monkey 2 it was 200 ms. Although the durations were different, the neural responses were integrated over the same 100 ms time window, which was delayed from the onset of the stimulus by 50 ms to account for the latency of the cortical response (see Figure 2). Also, recall that the median reaction time of the monkeys in the detection experiment was 200 ms (50 ms after the end of the 100 ms integration window).

**Figure 2.**
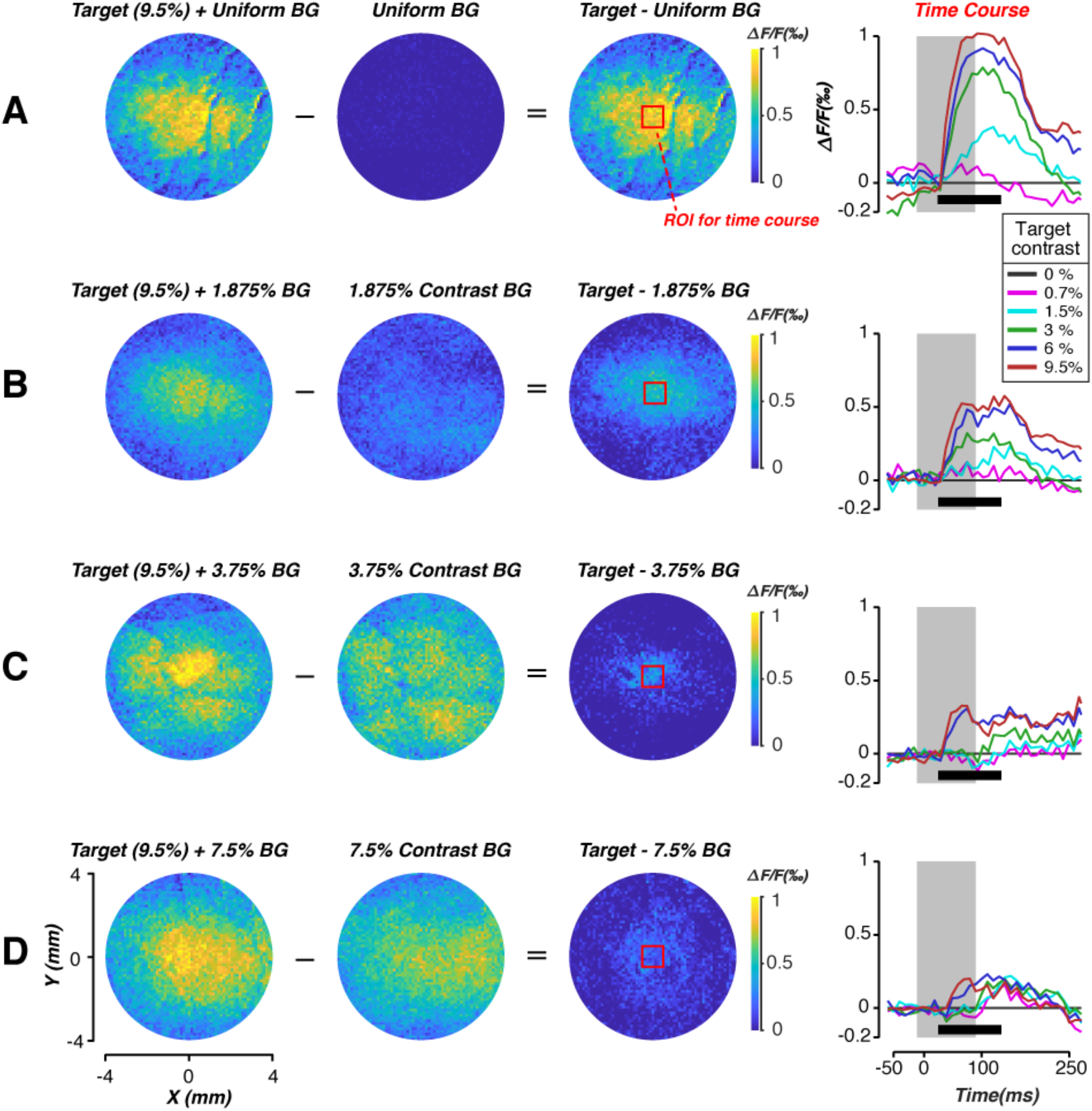
Example VSDI recordings from monkey 1. Each row (**A-D**) shows the average VSDI response for a particular background contrast (0%, 1.875%, 3.75%, 7.5% RMS), when the target is present and absent. The first two columns show the average responses when the target contrast is 9.5% RMS and 0% RMS (10 trials each), and the third column shows the difference in these average responses. The response images were derived by integrating for 100 ms beginning at response onset (which occurred at a fixed latency after stimulus onset). The last column shows the mean time courses in the central region outlined by the red box (1×1 mm^2^). Time courses were plotted after subtracting the mean time course of background-only trials. The grey rectangular shading indicates the stimulus presentation and the horizontal black bar indicates the integration interval (100 ms). Mean time courses are color-coded to represent the range of target contrasts used in the experiment (0%, 0.7%, 1.5%, 3%, 6%, and 9.5% RMS). Circular apertures are used for display purposes only.

### Estimating VSDI detection performance

Fifteen imaging sessions were collected from monkey 1, and ten sessions from monkey 2. The VSDI image frames were initially binned into 64 x 64 pixel image frames, where each pixel corresponded to 0.11 x 0.11 mm of cortex. Spatial binning filters out some of high spatial frequency shot noise in the camera images (Chen et al., 2006; Chen et al., 2012). Next, aberrant VSDI trials (< 5% of all trials) were identified and removed (see Chen et al., 2006). For each trial, we normalized (divided) the fluorescence amplitude at each pixel location by the average at that location measured during the first 100 ms of recording (which was before stimulus onset).

Following these preprocessing steps, neurometric functions were measured by applying a whitened template-matching (WTM) decoder to the population responses on each trial. The decoder was constructed in a way proposed by Chen et al., 2006. The WTM decoder is the optimal observer for detecting fixed targets in correlated additive Gaussian noise (e.g., see Brunelli & Poggio 1997), which is a good description of the noise in VSDI responses at the retinotopic scale (see Chen et al., 2006). In other words, our aim was to determine (approximately) the maximum discrimination performance possible given the information in the VSDI signals measured at the retinotopic scale. If down-stream circuits are using this information with a fixed efficiency, then we would expect neurometric performance to parallel behavioral performance.

To specify the WTM decoder for a given session, we first computed the average response in the 100-ms integration period (black horizontal bars in Figure 2), for each pixel location, when the background was uniform and the target was at its highest (9.5%) and lowest (0%) contrast (see Figure 2A). We then subtracted the 0% contrast image from the highest target contrast (9.5%) image. In general, we found that these incremental response profiles are approximately 2D Gaussians (Figure 3A). Thus, we fit the difference-response images from each session with a two-dimensional Gaussian function to obtain the “unwhitened” template. Figure 3B shows an example of this template.

**Figure 3.**
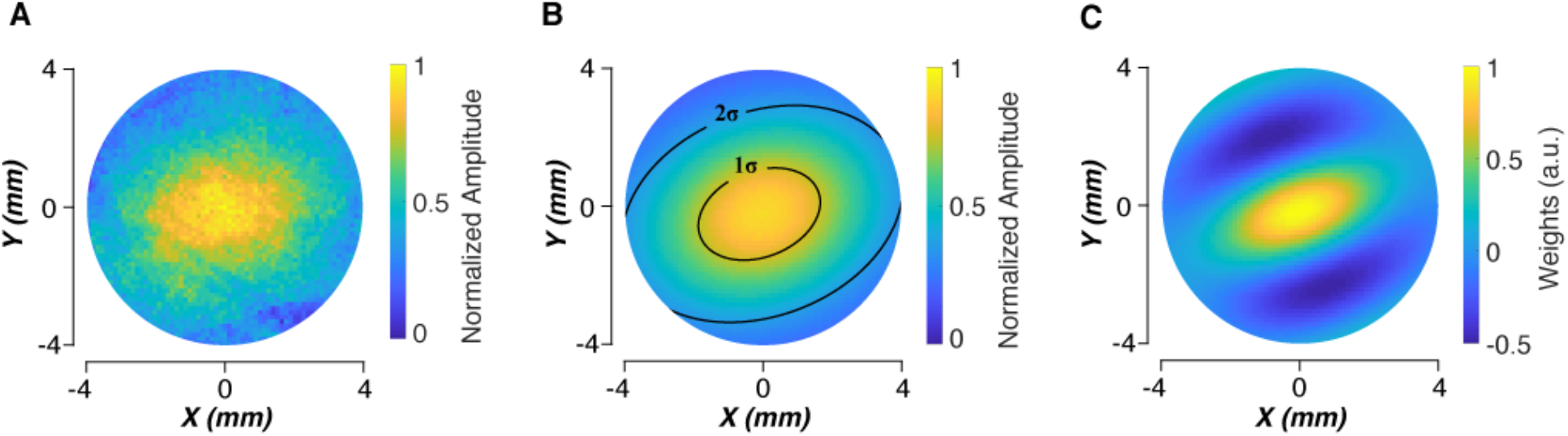
The whitened template matching (WTM) decoder. WTM decoders are approximately optimal for detection in correlated Gaussian noise, which is a good description of the noise in VSDI measurements. The WTM decoder applies a whitened template (weighting function) to the spatial response pattern on each trial and responds that the target is present if the template response exceeds a criterion. **(A)** The average difference in VSDI response (22 recording sessions: 14 sessions from monkey 1, 8 sessions from monkey 2) to a 9.5% contrast target and a 0% contrast target (target absent) on a uniform background. **(B)** The 2D Gaussian that was fit to the difference responses in panel A. For each experimental session, a 2D Gaussian was fit to the average difference response in the uniform background conditions. This fitted Gaussian was used in determining the whitened template for that session. **(C)** Example whitened template computed from panel B. The whitened template is a filtered version of the fitted 2D Gaussian that takes into account of the spatial noise correlations. Specifically, the antagonistic center-surround weights cancel much of the correlated noise, while removing little of the signal (see Chen et al., 2006 and text for more details). Circular apertures are used for display purposes only.

To determine the optimal “whitened” template we first measured the radially-averaged power spectrum of the VSD responses for the target-absent trials with a uniform background. We then divided the amplitude spectrum of the unwhitened template by the power spectrum (square of the amplitude spectrum) of the uniform-background signals to obtained the amplitude spectrum of the whitened template (Brunelli & Poggio 1997). Finally, we inverse Fourier transformed this amplitude spectrum to obtain the whitened template (e.g., Figure 3C; see Chen et al., 2006 for more details).

On each trial, the WTM decoder applies the whitened template to the population response (i.e., takes the dot product of the template and the population response) to obtain a single response scalar. If this response scalar exceeds a criterion the observer reports “target present”, otherwise “target absent”. Because we measure the real-valued response of the decoder, it is most efficient to estimate the detectability *d’* of the decoder by taking the difference in the mean responses divided by the square root of the average variance of the responses, and then convert to percent correct (see equation 3).

Notice that the whitened template is most positive in the center where the response to the target is largest and eventually becomes negative away from the center where the response to the target becomes negligible. To gain some intuition for why this whitened template is the near-optimal decoder we note that the noise correlations are substantial even at large distances from the center where the response to the target is weak or absent. This means that subtracting (negatively weighting) the responses at large distances cancels much of the correlated noise in the regions where the response to the target is strong, without cancelling much of the response to the target. This explains why the whitened template performs better than the simple unwhitened template (Chen et al., 2006).

In each experimental session, two neurometric functions were measured, one for the uniform background and one for natural backgrounds of some fixed contrast. We found that the neurometric functions for the uniform background varied substantially from session to session, almost surely because of variation of the quality of the dye staining and other non-neural scaling factors. Therefore, we combined the data across sessions using a simple model that accounts for the session-to-session variation in scaling factors.

Specifically, we assumed that the WTM response *r* on each trial is the sum of the neural response *r_n_* that is scaled by a dye efficiency constant for that session *a*_1_, plus a non-neural response *r*_0_ due to measurement noise:

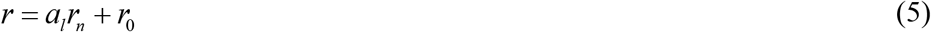

It follows that the detectability (*d’*) of the target is given by

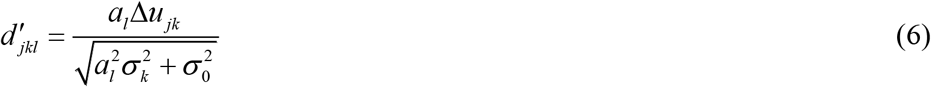

where *l* indexes the session, *k* indexes the background contrast, *j* indexes the target amplitude, *Δu_jk_* is the difference in the mean neural response to background plus target and background alone, *σ_k_* is the standard deviation of the neural response, and *σ*_0_ is the standard deviation of the measurement noise response. Furthermore, without loss of generality, we can absorb the standard deviation of the measurement noise and the standard deviation of the neural response for any one of the background contrast levels into the dye efficiency constants:

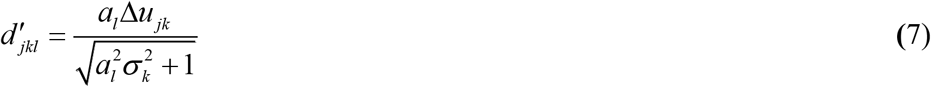

where *σ*_1_ = 1 (i.e., we absorb the standard deviation for contrast level 1 into the dye efficiency scalars). We used maximum likelihood methods to estimate simultaneously (from all data in all sessions for each monkey separately) the delta means, standard deviations, and the efficiency scalars. The estimated neural detectability for target amplitude *j* and background contrast *k* is then

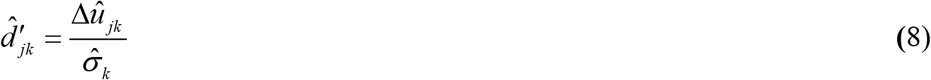

Note that when the value of *a_1_* is relatively low then the data from that session contributes relatively less (as it should) to the estimates of the delta means and standard deviations. We found that only one session, in one of the monkeys, had a really low value of *a_1_*. All other sessions contributed substantially to the estimates of detectability.

### Statistical analyses

Details of experimental procedures and visual stimuli are described above (see *Materials and Methods: Detection task* and *VSDI fixation task*). Behavioral performance was quantified by fitting psychometric curves using maximum-likelihood estimation. Summary statistics were derived separately for each individual subject, and standard errors were used to report across-subject variability. For VSDI responses, confidence intervals for the WTM decoder’s performance in each condition was estimated by bootstrap sampling (1000 iterations).

### Code/Software

Code is available upon reasonable request.

## Results

### Behavioral detection performance

Psychometric functions were measured in three monkeys for detection of a 4 cpd windowed sine-wave target presented at 2.5° eccentricity in randomly-selected natural backgrounds scaled to one of four different contrasts. Figure 4A shows the average bias-corrected psychometric functions for the three monkeys. The faint overlaid curves show the psychometric functions of the individual monkeys. As expected, the curves shift to the right as the background contrast increases. The gray circles in Figure 4C show the average thresholds corresponding to 69% correct (*d’* = 1; Note that the thresholds were obtained separately for each monkey and then averaged.) For comparison, the black circles show the average thresholds for three human observers for the same natural backgrounds at the same retinal eccentricity (but with a different range of background contrasts). The gray and black curves correspond to the generalized Weber’s law with the same slope and intercept parameters (equation 1) up to a single human-to-monkey scale factor *k_s_* = 2.5. The gray and black symbols in Figure 4D show the agreement after scaling. Clearly, monkey and human thresholds in natural backgrounds are consistent with a generalized Weber’s law, where the ratio of slope to intercept is the same for two species. This is evidence for a common neural computation and suggests that the macaque monkey is an appropriate animal model for human detection in natural backgrounds.

**Figure 4.**
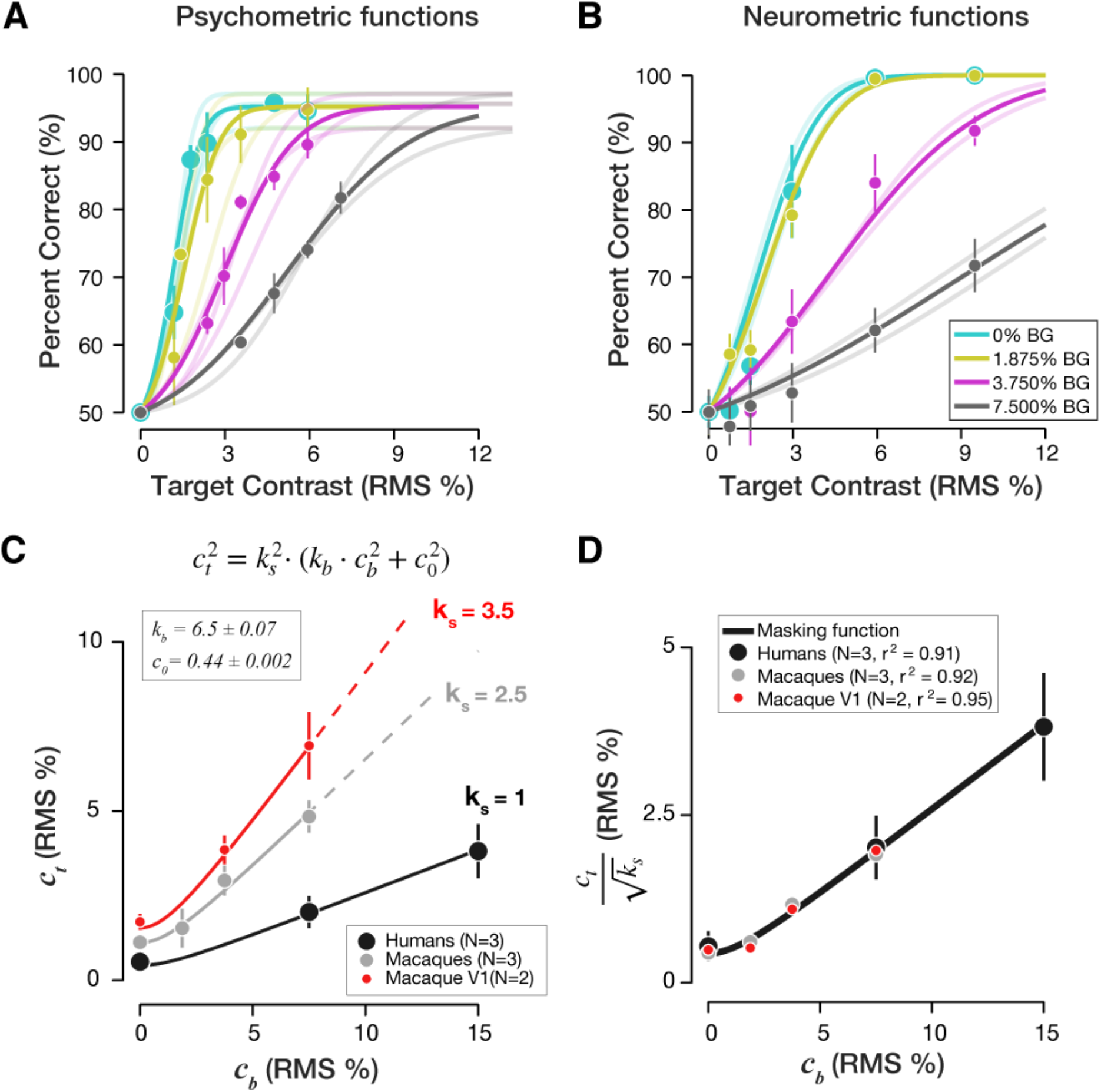
Behavioral and neural detection performance. **(A)** Bias-corrected psychometric functions of three monkeys for detection in uniform backgrounds and in natural backgrounds of several different RMS contrasts. The solid symbols and thick curves are the average psychometric functions. The thin curves are the psychometric functions of the individual monkeys. The error bars are standard errors across the three subjects. **(B)** Neurometric functions measured in two fixating monkeys. The solid symbols and thick curves are the average neurometric functions. The thin curves are the neurometric functions of the individual monkeys. The neurometric functions were obtained by applying a whitened-template-matching (WTM) decoder to voltage-sensitive-dye-imaging (VSDI) responses recorded at a retinotopic scale in primary visual cortex (V1). **(C)** Threshold as a function of background contrast. The gray symbols are the average behavioral thresholds of three monkeys. The black symbols are the average thresholds of three humans for the same targets and backgrounds (although with a different range of background contrasts; data from Bradley et al. 2014). The red symbols are the average neurometric thresholds of the WTM decoder applied to the VSD responses of two monkeys. The solid curves represent the generalized Weber’s law with the same slope and intercept parameters (*k_b_,c_0_*), but different overall scale factors (*k_s_*). **(D)** Agreement between thresholds of humans, monkeys, and V1 population responses. The black curve is the estimated generalized Weber’s law when *k_s_* = 1 (see the equation in C). The symbols are the thresholds of the humans, monkeys and V1 population responses after correcting for the effect of the scale factors.

### Neural detection performance

The consistency of macaque and human detection performance in natural backgrounds motivated us to make some initial measurements of the neural population responses in primary visual cortex of fixating macaque monkeys. These measurements were made in two additional monkeys that did not participate in the behavioral experiments. To assess the masking effects of natural backgrounds on detection sensitivity of neural population responses at the retinotopic scale in macaque V1, we applied a whitened template-matching (WTM) decoder (Fig. 3) to the single-trial responses. The appropriateness of this decoder is supported by the fact that the noise spectrum was similar (not shown here) across the contrast levels of the target and across the contrast levels of the natural backgrounds.

Figure 4B shows the average neurometric functions from the VSDI responses of the two monkeys for uniform and natural backgrounds of the same contrasts used in the behavioral experiments. The faint curves are the neurometric functions of the individual monkeys. In agreement with behavior, the neurometric functions of the WTM decoder shift to the right as the background contrast increases.

As in the behavioral experiments, we defined the neural threshold as the target contrast corresponding to *d’*=1. The red symbols in Figure 4C are the average neural thresholds for the two monkeys, and the solid curve shows the generalized Weber’s law for the exactly the same slope and intercept parameters as for the human and macaque behavioral data, but with a different overall scale factor (*k_s_* = 3.5). The red symbols in Figure 4D show the agreement after scaling.

The behavioral psychometric functions of the macaque monkeys (see Figure 4A) showed evidence of a small lapse rate of approximately 5%. We also computed neurometric functions assuming a 5% lapse rate in decision making. The scale factor for the neurometric thresholds increased slightly, but the agreement after scaling was as good as, or better than, that shown in

Figure 4D. We also varied the value of *d’* used to define threshold and found that the agreement after scaling was as good as that shown in Figure 4D. These results strengthen the evidence for a common neural computation and suggest that the macaque monkey (and human) behavioral thresholds are closely linked to the combined neural computations in retina and V1.

## Discussion

Natural selection guarantees that perceptual and cognitive mechanisms are relatively well matched to an organism’s natural tasks and stimuli. Thus, analyzing natural tasks and stimuli can be useful for obtaining principled hypotheses for neural computation. Furthermore, to fully understand neural computations it is crucial to measure and analyze behavioral and neural responses to the natural stimuli that the nervous system evolved to process. Here we measured psychometric and neurometric functions in the macaque monkey for detection of a simple target in uniform backgrounds and in natural backgrounds of various contrasts. We chose the macaque because anatomical, physiological, and behavioral studies (with simple stimuli) have shown the macaque visual system to be an excellent animal model of the human visual system.

We found that the psychometric functions and behavioral thresholds of three macaque monkeys measured at 2.5 deg eccentricity closely matched (up to a single overall scale factor) those of three human observers measured in an earlier study (Bradley et al. 2014) for the same stimuli at the same retinal eccentricity. Both humans and macaques followed the generalized Weber’s law as function of contrast for detection in natural backgrounds. Importantly, we found that the slope and intercept parameters of the generalized Weber’s law were the same in humans and macaques (up to a single scale factor) suggesting a common neural mechanism.

In a second study, we measured neurometric functions (for a subset of the same stimuli) from VSDI responses recorded at the retinotopic scale in the region of macaque V1 corresponding to ~2.5 deg eccentricity. The neurometric functions were computed using a whitened template matching (WTM) observer, which is near-optimal for VSDI responses at the retinotopic scale. We found that the neurometric thresholds measured in two macaques also followed the generalized Weber’s law with the same slope and intercept parameters (up to a single scale factor) as the behavioral thresholds of the three macaques and three humans. To analyze the data, we also introduced a simple new method for combining VSDI measurements across sessions that takes into account variation in the quality of dye staining and other non-neural scale factors.

The main conclusions are that (i) human and macaque ability to detect targets in natural backgrounds are very similarly affected by background contrast, (ii) these effects (generalized Weber’s law) suggest a common neural mechanism, (iii) population decoding at the retinotopic scale by down-stream circuits is consistent the behaviorally measured generalized Weber’s law, (iv) the macaque monkey is an appropriate animal model for gaining an understanding of the neural mechanisms in humans for detecting targets in natural backgrounds.

In the current study, which was based on Bradley et al. (2014), the natural background on each trial was adjusted to have a Gaussian gray-scale histogram so that every randomly selected background, even with an added high-contrast target, could be presented on a standard display without clipping. Also, the natural backgrounds were adjusted to have specific RMS contrasts. A different approach, based on constrained sampling from natural images, was used by Sebastian et al. (2017). In the constrained-sampling approach, millions of natural background patches the size of the target are sorted into a three-dimensional histogram along the three background dimensions known to have a big effect on detection performance: luminance, contrast, and structural (spatial frequency and orientation) similarity to the target. One advantage of the constrained-sampling approach is that no adjustments of the natural backgrounds are necessary because measurements can be limited to the space of bins where significant clipping does not occur. A second major advantage is that by sampling backgrounds from a sparse set of bins it is possible to measure, in modest-scale experiments, how the three dimensions affect detection performance individually and in combination. We also note that a limitation of the current study was that the VSDI measurements were made in a different set of fixating monkeys, and hence it was not possible to analyze the trial-by-trial correlations between the neural and behavioral responses. A logical next step, which we are currently pursuing, is to simultaneously measure VSDI responses and behavioral responses in constrained sampling experiments at both the retinotopic scale, and at the finer orientation-column scale where the neural effects of background similarity may become more apparent.

## Acknowledgements

This work was supported by US National Institutes of Health research grants EY024662, EY016454, and EY11747. The authors declare no competing financial interests.

## Notes

### Competing Interest Statement

The authors have declared no competing interest.

